# Peptidomics Mapping of Proteolysis Highlights Triple Activation of Sprouted Seeds by Germination, Homogenisation and Species Mixture

**DOI:** 10.64898/2026.01.08.698447

**Authors:** Indrani Bera, Raúl Fernández-Díaz, Michael O’Sullivan, Jean-Christophe Jacquier, Caitriona Scaife, Gabrielle Litovskich, Kieran Wynne, Denis C Shields

## Abstract

We investigated how seed proteolysis was enhanced by germination, by subsequent homogenisation (disrupting sprout compartments), and by co-incubation of homogenates from different species.

Mass spectrometry of released peptides tracked proteolytic signatures from chickpea, lentil, mung and broccoli proteins, in soaked seeds, in sprouted seeds, and after sprout homogenisation followed by incubation alone or in mixture with other sprouts. The proteolytic signatures differed markedly among the four species, and in the different treatment conditions. After homogenisation, legumain-like cleavage (after asparagine) increased in lentil, and proline-rich peptides increased in broccoli. For co-incubated homogenised sprouts, each species’ homogenate significantly contributed 6 to 57% of proteolytic patterns in peptides of other species, with chickpea and broccoli homogenates notably releasing metabolic protein peptides from mung and lentil.

Thus, germination, homogenisation and homogenate species mixtures can each contribute to proteolysis of seed peptides, potentially increasing digestibility and reducing allergenicity.

**HIGHLIGHTS:**

- Proteolysis motifs in soaked seeds and sprouts very diverse among species
- Seed germination proteolysis altered by homogenisation
- Seed germination proteolysis altered by co-incubation of different species
- Foods based on homogenised sprout mixtures may release more digestible peptides

## 1. INTRODUCTION

Proteolysis refers to the enzymatic degradation of proteins into peptides or free amino acids by proteases. This fundamental biological process plays an essential role in protein turnover, post-translational modification, and cellular signalling in all forms of life (Eckardt, 2024). In plants, proteolysis serves multiple physiological functions including seed germination, senescence, stress adaptation, and defence mechanisms (van der Hoorn, 2008). Proteolytic cleavage is essential for the maturation and breakdown of storage proteins in seeds. In this study, we wanted to explore how the activities and impacts of various proteases could be tracked during germination of various seeds, and how their activities might be altered not only by germination, but also by homogenising sprouts and by mixing preparations from different sprout species.

Plant protease families are serine, cysteine, aspartic, metalloproteases and threonine proteases, each named after a key residue or feature of their catalytic sites (Dunn, 2001; Turk et al., 2006; Figueiredo et al, 2021; Grudkowska & Zagdanska, 2004). Cysteine endoproteases play a key role in breaking down seed storage proteins (Grudkowska & Zagdanska, 2004). These include papain-like vignains (often cleaving after R or K), and legumains (typically cleaving after N in plants, Rotari et al,2001). Legumains control proteolysis of seed storage proteins during maturation and packing, as well as in further proteolysis during germination (Grudkowska & Zagdanska, 2004). A single calcium dependent cysteine protease, calpain, has roles in seed development, and its substrate specificity is not clearly defined by a dominating amino acid preference (Margis et al, 2003; Liu et al,2011). Caspases cleaving after D, are found in the cytosol (Korthout et al, 2000), in contrast to legumains (pH optimum 6.0 , Rotari et al,2001) and vignains, which are both typically located in the storage vacuoles where the protein bodies of seed storage proteins accumulate.

Aspartic endoproteases (Gruis et al, 2002) include barley phytepsin, which is stored in the protein bodies of ungerminated seeds, also play a role in storage protein breakdown (Marttila et al, 1995). There is limited evidence for a role for a serine endopeptidase cleaving D-rich regions being expressed at day 6 of soya bean germination (Liu et al 2011).

Serine carboxypeptidases are present during monocot germination (Washio and Ishikawa, 1994) and in legumes, where phaseolain has an apparent preference for cleavage after I, L and V (Carey and Wells, 1972). Other exopeptidases are likely to also play key roles, with a proteomics characterisation of an extract based on soaked sesame seeds identifying likely exopeptidases at abundances often equivalent to that of the major endoproteases vignain and legumain, identifying in order of abundance tripeptidyl-peptidase 2, leucine aminopeptidase 2, proline iminopeptidase, serine carboxypeptidase-like, cytosolic oligopeptidase A, aminopeptidase M1, Xaa-Pro aminopeptidase P, puromycin-sensitive aminopeptidase, aspartyl aminopeptidase, serine carboxypeptidase-like 42, and glutamate carboxypeptidase (Chen et al, 2021). Methionine aminopeptidase and lysosomal Pro-X carboxypeptidase are detectable in germinating chickpeas (Bera et al 2024), and there are potential distinct exopeptidases removing amino-terminal residues; aminopeptidase M1 [L/P/Y] (Murphy et al, 2002), aminopeptidase P [X|P] (Hauser et al, 2021), methionine aminopeptidase [M] (Jeong et al, 2011), and carboxy-terminal residues; serine carboxypeptidase [ILV] (Zuber and Matile, 1968), glutamate carboxypeptidase [E] (Shi et al 2013) and prolyl carboxypeptidase [P|X] (Khaket et al ,2015) .

Not only is there a diversity of endo- and exoproteolytic potential activities, but they appear to vary substantially over the course of germination (Bera et al 2024). The germination process is initiated by imbibition and involves the release of gibberellins from the embryo, which stimulate the synthesis and secretion of hydrolytic enzymes including proteases from the aleurone layer (Nonogaki et al., 2010). The above proteases come together to play roles in germinating seeds, including auto-activation and activating other proteases, breaking down protease inhibitors and proteases, and breaking down the seed storage proteins to release carbon and nitrogen reserves to the growing sprout. Regulation of these processes includes multiple factors, such as the change towards a more aqueous environment, activation of pre-translated proteases present in the seed, differential expression of protease gene families, and dynamic interactions between proteases and storage proteins across various seed compartments (Muntz, 1996).

Seed germination shows a dynamic change in bulk protease cleavage over time and in a strain-specific manner, with different proteases coming to the fore at different stages, as seen from endoprotease preferences identified by mass spectrometry of released peptides in chickpea (Bera et al, 2023; Nitride et al, 2022). These changes may be controlled in part by pH, substrate-enzyme access, removal of protease auto-inhibitory domains, and colocalisation with protease inhibitors. An example of the latter is the barley cystatin inhibitors whose QxVxG motif interacts with legumains (Martinez et al, 2009). All these components are very incompletely mapped, making it hard to draw biological inferences from the co-expression network of proteases and inhibitors, as has been investigated in human intracellular cell signalling (Fortelny et al, 2014). In chickpea, legumain-like and vignain-like proteases are activated at distinct stages of germination, each recognizing specific peptide motifs (Bera et al., 2024). These stage-specific proteolytic patterns suggest a complex orchestration of enzymatic activity which is likely to be aligned with metabolic and other needs of the developing seedling, such as host defence. Thus, germination enhances legume seed protein hydrolysis by reducing protease inhibitors, activating proteases, and breaking down storage proteins. Understanding these changes can guide breeding and processing methods to improve nutritional quality (Bera et al., 2023). In a comparative study across legumes, *in vitro* protein digestibility (IVPD) increased during germination, with notable enhancement by day five in chickpeas and other species (Uppal et al., 2012).

Food processing techniques involving homogenisation and addition of external proteases can serve to increase proteolysis. High pressure homogenization techniques can disrupt cellular compartments, thereby increasing enzyme-substrate interactions, and by changing the secondary structure and physicochemical properties of seed proteins (Liang et al., 2022; Melchior et al., 2022), with implications for food texture, flavor, and nutritional value. External proteases can modify the digestibility, functionality, or allergenicity of plant proteins. Fungal proteases from *Aspergillus oryzae* and 2 other species have demonstrated high efficacy in hydrolyzing legume (pea) proteins and increasing *in vitro* protein digestibility (Yadav et al., 2022). The plant-derived protease bromelain has been used to improve the antioxidant properties and anti-inflammatory properties of legume protein (Xu et al., 2021). Papain and bromelain hydrolysis of chickpea and lentil released small (200 to 1000 Da) peptides with antioxidant and antihypertensive properties. These applications highlight the potential use of cross-species proteases in activating bioactive health-promoting food components (Domokos-Szabolcsy et al., 2015).

Here, as an alternative to addition of externally processed enzymes and harsh treatment conditions, we wished to explore the potential of gentler food processing steps (seed germination, room temperature homogenisation and species mixtures) to alter the signatures of proteolysis of seed proteins. We used LC-MS/MS peptidomics to explore how sequential events of germination, homogenisation and cross-species co-incubation of seeds can alter proteolysis in sprouts of the legumes chickpea, lentil and mung, and of the brassica broccoli. We found that each of these processes had considerable effects on the proteolysis of seed sprout proteins, with a marked diversity of proteolytic signatures across species and conditions.

## 2. MATERIALS AND METHODS

### 2.1 Materials

Diethyl ether (99.5% purity), NaCl (>99.5% purity), Pierce Trypsin Mass Spectrometry grade, Iodoacetamide ultra-pure (98% purity) proteomics Grade, TCEP (98% purity), Pierce Water LC-MS Grade, Acetonitrile (99.9%), Trifluoroacetic acid (99.5%) biochemistry grade were procured from VWR chemicals and 0.1% (v/v) formic acid LC-MS grade was procured from Fisher Scientific.

### 2.1 Preparation of sprouts and sprout mixtures for peptide extraction

Chickpeas, lentil, mung and broccoli seeds were obtained from Happy Pear Ltd, Greystones, Ireland. They were germinated as biological replicates in three separate weeks, following a protocol consistent with commercial sprout production. Germination is considered to be finished when the radicle comes out from the seed coat (Bewly and Black, 1994). Accordingly, chickpea and broccoli were germinated for 4 days while mung and lentil were germinated for 2 days. The seeds were soaked for 8 hours and a third of the soaked seeds was taken out and ground using a food grade homogenizer and stored at -20°C. These represent the Day 0 of germination. The remaining two-thirds were then germinated at 18°C while washing them twice daily to avoid drying out or overheating. The excess moisture was dried out after every wash. After germination was finished, the sprouts were weighed and divided into 6 parts. One part was ground and stored at -20°C immediately, which represents the germinated samples (4-day sample for chickpea and broccoli and 2-day sample for mung and lentil). Another part was ground and kept incubated at room temperature (20°C) which represented a germinated+4 hours (incubated) sample. The remaining fractions were homogenized in a pairwise fashion with the other sprouts and incubated for 4 hours at room temperature.

### 2.2 Peptide extraction and filtration

Sprout mixtures from all time points as discussed above (with 3 replicates each) were resuspended as separate aliquots by dissolving 5g of sprout mixture homogenate in 0.2M NaCl solution in glass beakers as 10% w/w solutions, bringing the total volume to 50 ml. The dispersions were stirred for 2 hours by a magnetic stirrer at room temperature and then placed in the refrigerator overnight. Next day, the extracts containing soluble peptides were collected by centrifuging them in polypropylene tubes at 12,000g for 20 minutes at 20°C. The soluble supernatant (15 mL) combined from NaCl and water extraction was transferred very carefully into a 10kD molecular weight cut-off centrifugal filtration module (Amicon Ultra Centrifugal Filter, 10 kDa MWCO) and centrifuged at 5000g for 20 minutes. The collected permeate fractions were stored at -20°C before injection into the liquid chromatography (LC) column for LC/MS-MS. Two technical replicates of the mass spectrometry analysis were obtained from each of the biological replicates.

### 2.3 Peptide detection using LC-MS/MS

Proteomics samples were loaded onto EvoTips and run on a timsTOF Pro mass spectrometer (Bruker Daltonics, Bremen, Germany) coupled to the EvoSep One system (EvoSep BioSystems, Odense, Denmark). The peptides were separated on a reversed-phase C 18 Endurance column (15cm x 150μm ID, C 18, 1.9 μm) using the preset 30 SPD method. Mobile phases were 0.1% (v/v) formic acid in water (phase A) and 0.1% (v/v) formic acid in acetonitrile (phase B). The peptides were separated by an increasing gradient of mobile phase B for 44 minutes using a flow rate of 0.5 uL/min. The timsTOF Pro mass spectrometer was operated in positive ion polarity with TIMS (Trapped Ion Mobility Spectrometry) and PASEF (Parallel Accumulation Serial Fragmentation) modes enabled. The accumulation and ramp times for the TIMS were both set to 100 ms., with an ion mobility (1/k0) range from 0.6 to 1.6 Vs/cm. Spectra were recorded in the mass range from 100 to 1,700 m/z. The precursor (MS) Intensity Threshold was set to 2,500 and the precursor Target Intensity set to 20,000. Each PASEF cycle consisted of one MS ramp for precursor detection followed by 10 PASEF MS/MS ramps, with a total cycle time of 1.17 s. The mass spectrometry data have been deposited to the PRIDE Archive (http://www.ebi.ac.uk/pride/archive/) via the PRIDE partner repository (Project accession: PXD072115, Username: Password: 77RvUcbwWSmu).

### 2.4 Analysis of LC-MS/MS data

Raw mass spectrometry data from 19 runs (saved in the Bruker formatted .d directory) were analyzed using the FragPipe software suite (Hsiao et al, 2024) (v20), integrating MSFragger (v3.8) for peptide identification, IonQuant (v1.9.8) for quantification, Philosopher (v5) for spectral validation and protein inference, and Python (v3.9.7) for downstream processing. The workflow was configured for a non-specific peptidome search with MS data type set to IM-MS. Decoys were appended to the FASTA databases for target-decoy-based FDR estimation. MSFragger was run with a peptide length range of 7–65 amino acids. For label-free quantification (LFQ), IonQuant (v1.9.8) was used for MS1 quantification mode within FragPipe. The MaxLFQ algorithm was utilised with the minimum ion requirement set to 1, allowing for the inclusion of single ion observations in the quantitation process. This parameter setting was chosen to maximise the sensitivity of the LFQ analysis, enabling the detection and quantification of low-abundance peptides within the sample matrix. Species-specific protein databases were downloaded from UniProt (on 08/01/2024) for each sample type: Chickpea samples were searched against *Cicer arietinum* (taxonomy ID: 3827, number of proteins: 48,501); and Mung bean samples were searched against *Vigna radiata* (taxonomy ID: 157791, number of proteins: 49,280). In the absence of a sequenced lentil proteome available, lentil samples were searched against the *Pisum sativum* UniProt database (taxonomy ID: 3888, number of proteins: 66,521 proteins) and *Lens culinaris* (taxonomy ID: 3864, number of proteins: 888 proteins) due to evolutionary proximity (Choi et al., 2004); Broccoli samples were searched against *Brassica oleracea* (taxonomy ID: 3712, number of proteins:121,738 proteins) (Guo et al., 2021). Default parameters were used unless otherwise specified, and all searches were performed within the FragPipe interface. The workflow was run on 20 CPU cores with 2 threads per core. Contaminants and reverse sequence decoys were corrected using Philosopher. Subsequent data analysis utilised the protein and peptide “.tsv” files after post-processing.

### 2.5 Proteolytic Motif Extraction and Analysis

Quantitative analyses were carried out using R (R Core Team, 2013), using the Biostrings package to import the plant FASTA files. Peptide-spectrum match (PSM) data obtained from FragPipe output files were read and merged with the corresponding protein sequences based on UNIPROT protein IDs. N-terminal and C-terminal cleavage motifs (±4 residues surrounding the cleavage site, corresponding to the P4-P1 residues upstream, and P1’-P4’ sites downstream) were extracted from full-length protein sequences for each peptide, and only unambiguous motifs of exact length (8 amino acids) were aggregated to define a motif profile for each sample type and terminus type. Motif profiles were one-hot encoded into binary vectors for each amino acid at the eight positions. Our analysis was restricted to the 100 proteins with the highest count of peptides. Sequences of the 100 proteins with the highest peptide counts were used to generate a background distribution of amino acid frequencies for motif enrichment analysis. A binomial probability model (O’Shea et al , 2013) was implemented in R assuming *a priori* the same null background distribution of amino acids at each cleavage position; this estimated the statistical significance of amino acid enrichment or depletion at each cleavage site position, and the values were represented as a stacked sequence logo with the most significant amino acids having the largest letter height.

### 2.6 Regression quantification of inter-species proteolytic effects in sprout mixtures

Given the proteolysis profile for two sprout species incubated separately, it is possible to ask whether the co-incubated sprouts show a marked effect of the proteolysis effects of each species acting on the peptides of the other species. If we assume the effects are linear and additive, the contributions of each species to the cleavage of the set of peptides from each species may be estimated, with confidence intervals, by linear regression. Across the eight motif positions, we defined the frequency of amino acid *R*_*i*_ in position *P*_*j*_ as the number of peptides with *R*_*i*_ in position *P*_*j*_divided by the total number of peptides. Assuming that these frequency measures are a realistic approximation of the underlying probability distributions, these data comprise the probability position matrix (PPM) describing how probable it is that each amino acid appears at each position across all the sequences. Each PPM consisted of 8 columns representing cleavage positions (P4–P4’) and 20 rows indicating the proportion of each amino acid residue. Contributions of the two isolated incubated PPMs to the co-incubated (mixed) PPM were estimated according to the following equation:

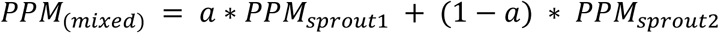

where *a* represents the relative contribution of sprout1, and *(1-a)* represents the contribution of sprout2 and *mixed* = *sprout*1 + *sprout*2. Thus, we can model the mixed proteolytic pattern (*PPM*_*mixed*_) as the sum of the proteolytic patterns of each sprout in isolation (*PPM*_*sprout*1_ and *PPM*_*sprout*2_). This model was fitted using Linear Least Squares regression, obtaining estimates of the regression coefficient *a* and its 95% confidence intervals. If 0 is within the confidence interval, the contribution of *PPM*_*sprout*1_ is negligible; if 1 lies within the confidence interval, then the contribution of *PPM*_*sprout*2_ is not statistically significant.

## 3. RESULTS AND DISCUSSION

To explore proteolytic activities in sprouting seeds, we performed a comparative peptidomic analysis of the legumes chickpea, lentil and mung, along with broccoli as a non-legume for comparison. We contrasted soaked seeds with germinated sprouts, sprouts before and after homogenisation/incubation, and sprout homogenates co-incubated together. Peptides under 10 kD were cleaned up for each sample, and identified using LC-MS/MS. We focussed on analysis of the peptides found in the top 100 proteins, to maximise the focus on proteolysis of the major seed storage proteins. The overall study design is visualised in Figure 1.

**Figure 1.**
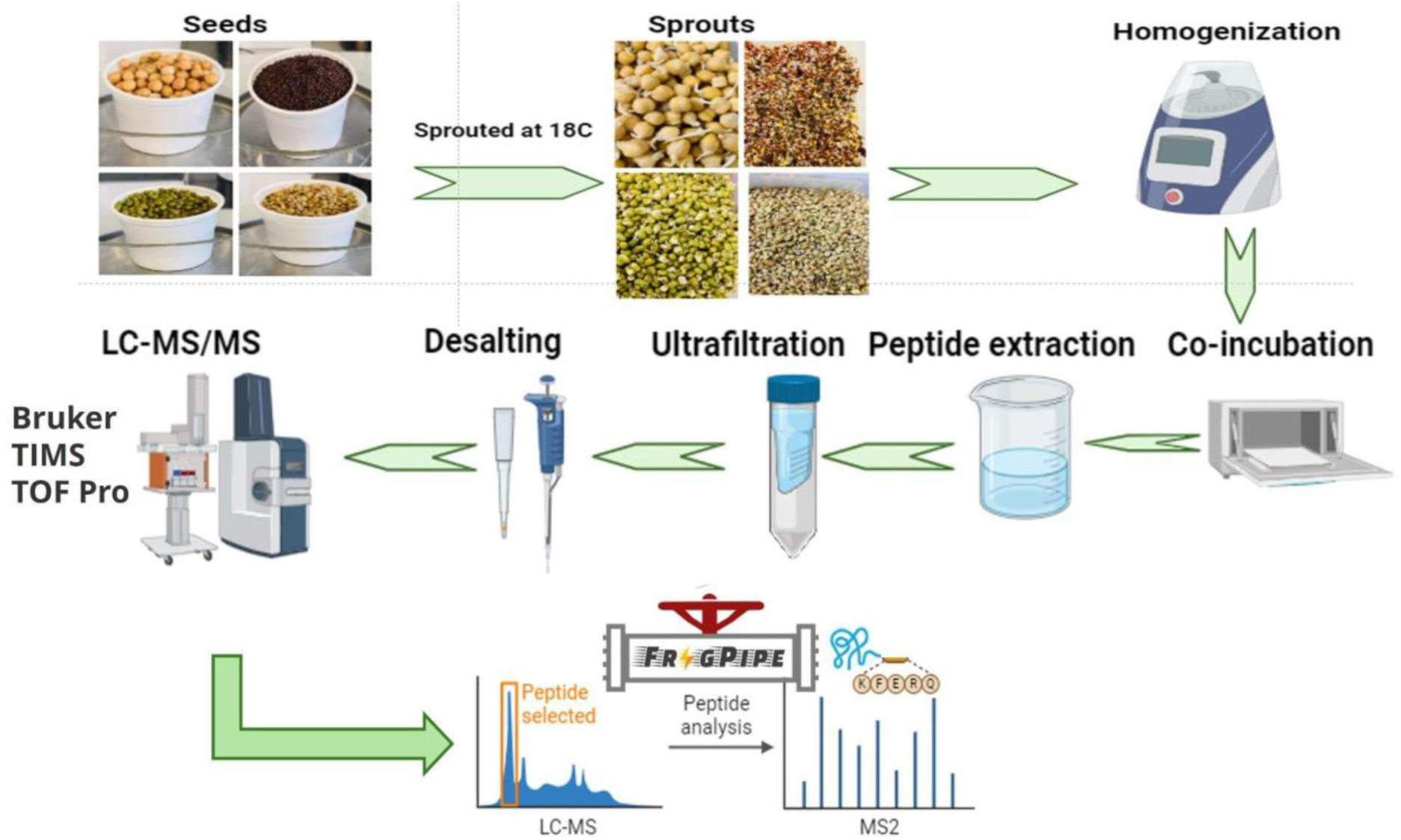
Overall workflow to identify sprout cooperative proteolysis.

Table 1 shows the number of peptides and proteins identified from each sample. The number of peptides identified from lentil were lower, likely reflecting the use of the closely related pea *Pisum sativum* as the search database, in the absence of a proteome derived from genome sequencing of lentil. Germination increased the number of detected peptides in chickpea and mung, while homogenisation increased further in all four species. In the coincubated homogenates of species pairs, there were reasonable numbers of peptides from both contributing species, allowing us to characterize below the shifts in the proteolytic profiles induced by the introduction of components from the other species.

**Table 1.** Number of peptides and proteins identified across three biological replicates for each sample. For species mixtures, the breakdown of the peptide count into those assigned to the two species are shown in parentheses, in the same order as in the Species column.

| Species | Time after soaking | Peptide count | Protein count |
| --- | --- | --- | --- |
| Chickpea | 0 day | 2340 | 203 |
| Chickpea | 4 days | 8251 | 543 |
| Chickpea | 4 days+ 4h incubated homogenate | 13142 | 953 |
| Broccoli | 0 day | 2982 | 669 |
| Broccoli | 4 days | 2051 | 408 |
| Broccoli | 4 days + 4h incubated homogenate | 7601 | 1011 |
| Lentil | 0 day | 253 | 69 |
| Lentil | 2 days | 254 | 72 |
| Lentil | 2 days + 4h incubated homogenate | 2926 | 336 |
| Mung | 0 day | 779 | 133 |
| Mung | 2 days | 3575 | 384 |
| Mung | 2 days + 4h incubated homogenate | 4090 | 530 |
| Chickpea + Broccoli | Combined incubated homogenate | 8492 [6922+ 1570] | 826 [411+415] |
| Chickpea + Mung | Combined incubated homogenate | 8180 [5607+2573] | 868[536 + 332] |
| Chickpea + Lentil | Combined incubated homogenate | 7038 [6030 + 1018] | 759 [489 + 270] |
| Lentil + Mung | Combined incubated homogenate | 2264 [722 + 1542] | 328 [132 + 196] |
| Lentil + Broccoli | Combined incubated homogenate | 2860 [1491 + 1369] | 410 [164 + 246] |
| Mung + Broccoli | Combined incubated homogenate | 4494 [3014 + 1448] | 700 [326 + 374] |
| Mung+Lentil+<br>Broccoli | Combined incubated homogenate | 4627 [1629 + 546 +<br>2452] | 761[189<br>+128+444] |

### 3.1 Distinct patterns at N and C termini indicate variable exoprotease efficiencies

Protease preferences may be both positive and negative, and the measure of statistical enrichment used here is also used to provide a measure of statistical avoidance, represented as amino acids below the line in the sequence logo. While some avoidances were seen (particularly P1-[VI], where square brackets group the alternative residues at the same position), these did not show such marked differences between species and mixtures, and accordingly we focussed on the enriched amino acids. The specificities of the three main protease types (endopeptidases, carboxypeptidases and aminopeptidases) will each contribute to the distribution of peptides seen in a sample. Protease substrates are typically characterised (e.g. Figure 2) by the the four residues upstream of cleavage (P4 to P1), and the four downstream (P1’ to P4’), with the main specificities of plant proteases residing at the P1 position. Under the simplifying assumption that upstream and downstream regions of cleavage sites are independent, then the P4 to P1 specificities are best estimated from the N-terminal cleavage sites, since marked biases in N-terminal degradation of peptides by aminopeptidases will not affect the upstream sequences as strongly. Similarly the P1’ to P4’ specificities are best explained by the carboxy-terminal cleavage motifs. Exopeptidase biases that influence endopeptidase patterns can be summarised as three possible main effects: (1) rapid elimination of endopeptidase motifs from termini by proteases that are more efficient for those motifs, (2) accumulation of motifs that are resistant to exoprotease activity, and (3) a possible “smearing” of the motif across adjacent termini by variable extents of digestion from a common endoproteolytic terminus. We did not find evidence of the latter effect in the data, suggesting that detectable peptides might be rapidly removed from the detectable size range, by additional endoproteolysis or by rapid sequential exoproteolysis. We did find evidence of the first two effects, described below. Given these biases, we focussed on the N-terminal peptide apparent cleavage sites as the best reflection of endoproteolysis P4 to P1 preferences, and the C-terminal cleavage sites as a reflection of P1’ to P4’ preferences.

**Figure 2.**
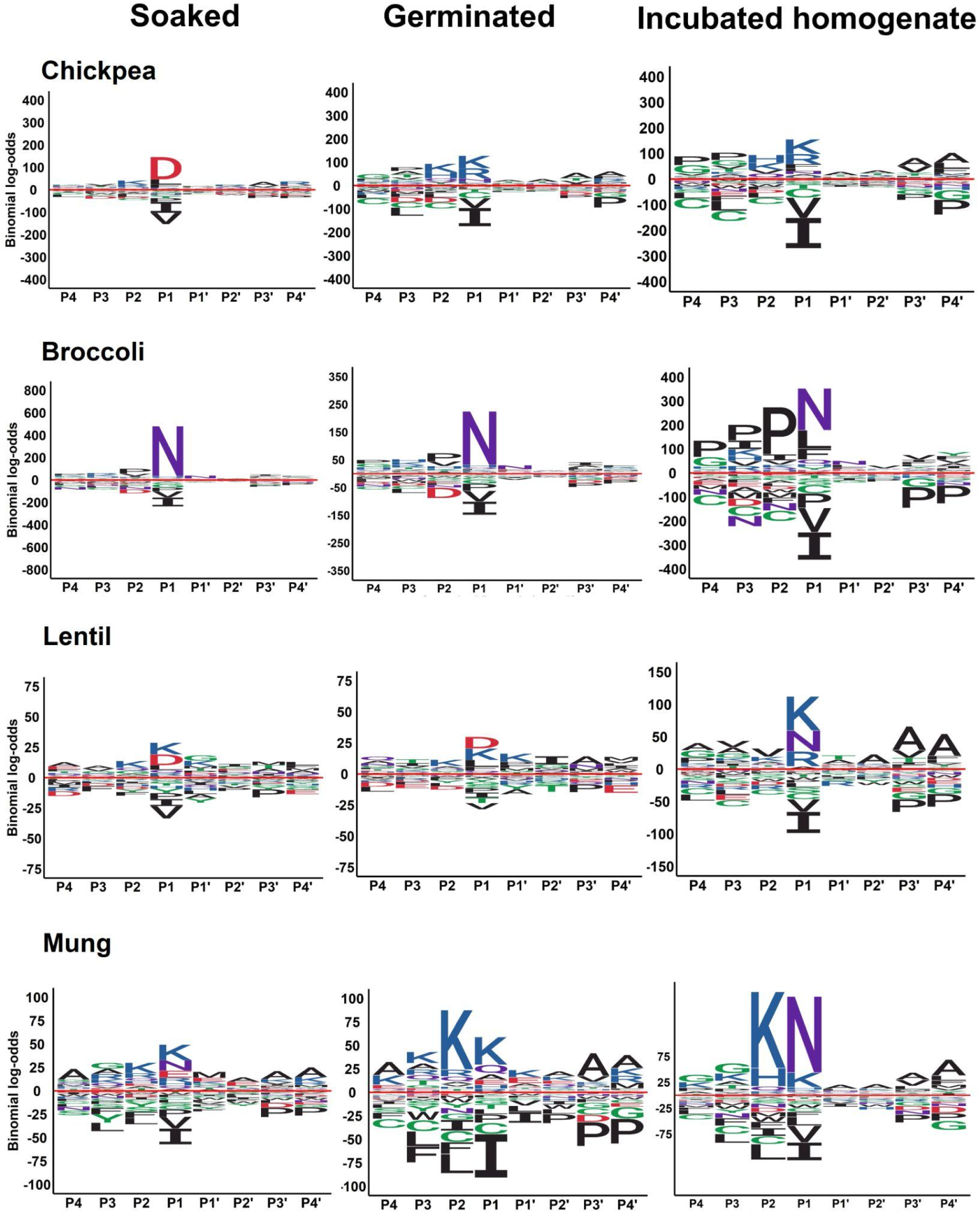
C-terminal proteolytic motifs of Chickpea, Broccoli, Mung and Lentil. Height of letter is proportional to statistical significance of enrichment.

### 3.2 Distinctive patterns between species of soaked seeds and sprouts

The four species showed distinct profiles over the soaked and germinated samples (Figures 2 and 3). Broccoli and lentil showed little shift in profile between soaked and germinated seeds (as seen for soaked versus 2-day germinated chickpea, Bera et al 2024), while chickpea and mung showed a stronger shift in profile after germination. Differences in the chosen numbers of days post germination in this study reflected potential harvesting times for use of different sprouts. Among the P4 to P1 sites (Figure 3) P1 conferred greatest specificity, while among the P1’ to P4’ sites interpreted from the C-terminal dataset, more modest effects were observed at P3’ and P4’ (Figure 2).

**Figure 3.**
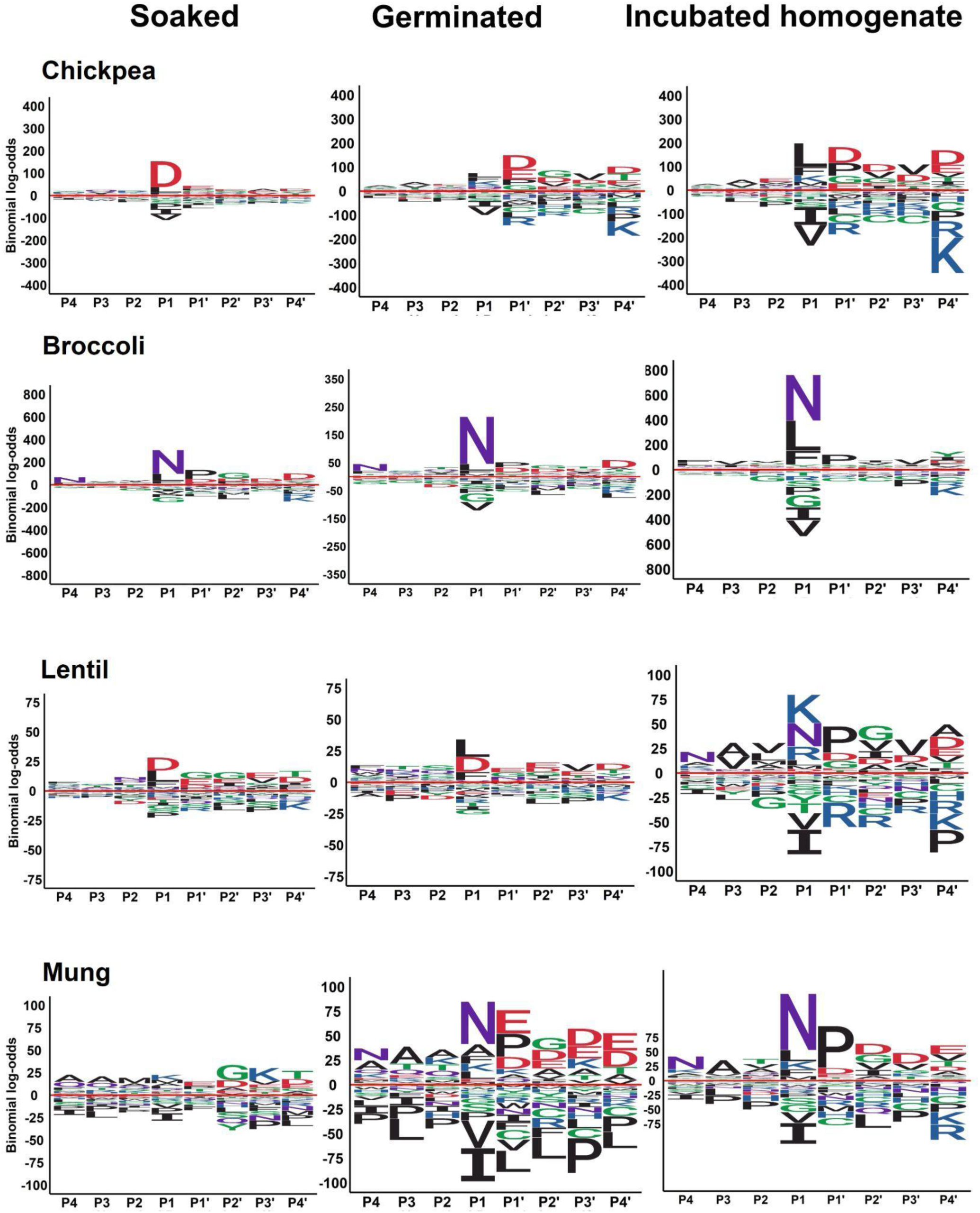
N-terminal proteolytic motif of Chickpea, Broccoli, Mung and Lentil.Height of letter is proportional to statistical significance of enrichment.

In soaked seeds, chickpea and lentil showed similar P1-[DL] preferences with a different preference in broccoli (P1-[NL]) and mung (no strong preferences). In germinated seeds, P1-N was preferred in broccoli and mung, indicating an up-regulation of legumain activity (Bera et al 2024) in mung. No such enrichment was seen in chickpea and lentil soaked or germinated seeds (although P1-N enrichment was seen at other timepoints for chickpea in a previous study, Bera et al 2024).

P1L was most strongly preferred in lentils, with variable levels of enrichment also seen in broccoli, mung and chickpea, similar between soaked and germinated seeds in all four species (Figure 3). The preference for P1-[LF] may reflect a plant phytepsin-like aspartic endopeptidase cleavage after L and F (Chen et al, 2021, Kervinen et al, 2004), and representing 35 of 62 cleavages documented on the MEROPS database ( Phytepsin A01.020, Kervinen et al 2004). Phytepsins have been found in large quantities in germinating seeds. Purified phytepsins can break down seed storage proteins *in vitro* in a variety of species (Kervinen et al, 2004), including in Euphorbiaeae vacuoles (Hiraiwa et al 1997), and wheat gluten after aspartic acid at low pH (Bleukx et al 1998). In mung bean, a short window of activity of the aspartic endoprotease VrAP peaking shortly after soaking is negatively correlated with the activity of an inhibitor, suggesting it may play a highly controlled role in mung bean germination (Kulkarni and Rao,2009).

P1-D may reflect caspase-related metacaspase activity, which is involved in seed quiescence (Julien and Wells, 2017), activating seed dormancy by accumulating storage proteins (Liu et al, 2024). This pattern was not seen (Bera et al 2024) in chickpea soaked for a longer period (12 hours versus 8), suggesting a short time period over which P1-D can be detected in chickpea. Here, it was absent in germinated chickpea but persisted in germinated lentil (Figure 3). Chickpea shows a modest preference for P1-K after germination (Figure 3), reflecting a likely vignain-like activity (Bera et al, 2024).

### 3.3 Exoproteolytic effects and the differences between amino and carboxy termini

Previous datasets (Bera et al 2024, Nitride et al ref) implicate a set of digestion resistant peptides associated with a distinct profile of residue enrichment at amino termini of chickpea peptides; these peptides appear resistant to proteolysis during bo germination, and during human gut digestion. The principal feature of these peptides is enrichment for the negatively charged amino acids aspartic acid (D) and glutamic acid (E) at the first four residues of the peptide (P1’ to P4’ sites). This effect is clearly seen again in this study for chickpea, as a set of enriched negative amino acids in the peptide itself at the amino terminus (compared with the carboxy terminus, Figs. 2 and 3), not only in the soaked seeds, but increased after 4 days sprouting. This appears consistent with protease resistant peptides accumulating over time, as previously hypothesised (Bera et al 2024). Interestingly, a very similar enrichment is seen for mung and lentil. The effect is present in broccoli but is weaker. Thus, these peptides appear resistant to aminopeptidase or further endoproteolytic cleavage, and accumulate. Other patterns like this, such as the P2’-G enrichment in mung that is only seen in the amino-terminal cleavage sites (Figs. 2 and 3), likely reflect a similar resistance to aminopeptidase breakdown. At the C-terminus, the enrichment of P2-K in mung (and to a lesser extent in chickpea and lentil) could reflect carboxypeptidase resistance.

Notably, some amino terminal P1 preferences were not seen in the carboxy termini of soaked and germinated seeds, in particular [LF] in germinated broccoli and chickpea (Figure 2 and 3). This could well reflect rapid elimination after phytepsin-like endoproteolysis by a specific carboxypeptidase, such as phaseolin (Carey and Wells, 1972). Similarly, the disappearance from germinated C-termini of the mung P1-N seen at N-termini could possibly reflect specificity of an unidentified carboxypeptidase.

### 3.4 Homogenisation of sprouts alters the proteolytic signatures

Homogenisation of sprouts followed by four hours of incubation at room temperature has the potential to release new proteolytic activities on substrates, as proteases and substrates from different seed and subcellular compartments come into contact with each other. Pronounced shifts in P1 preferences were seen (Figure 3). Chickpea and broccoli showed an increase in the pre-existing P1-[LF] enrichment after homogenisation (Figure 3), but this activity showed a marked decline in lentil (Figure 3); this reduction of a phytepsin-like protease signature may reflect down-regulation of the phytepsin rather than a marked increase in phaseolin-like carboxypeptidase removal of the signature, since it is also lower in Figure 2. Lentil also lost any caspase-like P1-D enrichment, with an upregulation of vignain-like (P1-[KR]) and legumain-like (P1-N) preferences.

P1’-P4’ changes in preferences on homogenisation (Figure 2) show a marked increase in P3’-A and P4’-A preferences. These appear to increase across various legume samples whenever the vignain-like P1-[KR] are increased (Figure 3), suggesting that it may be a component of the vignain-like preference in legumes. The variation across different legume samples (see also Bera et al 2024) suggest that the various vignains seen within each legume are likely to have different P1 specificities for K and R.

Homogenisation has the potential to break down accumulating less digestible peptides. On homogenisation, there appears to be some reduction of the negatively charged motifs seen at peptide N-termini in broccoli and mung, but not much reduction in chickpea (Figure 3). Homogenisation appears to increase broccoli peptides rich in proline near their C-termini, and legume peptides rich in G and K near their C-termini, with the most striking preference seen for P2-K in mung (Figure 2). Homogenisation also increases the frequency of N-terminal prolines in all four species (Figure 3), suggesting that these peptides are being released more often by endoproteolysis or are becoming harder to eliminate (e.g. if homogenisation down-regulates prolyl aminopeptidase activity), and so they accumulate in the samples.

### 3.5 Changes in proteolytic profiles induced by co-incubation with other sprouts

We sought to understand if the proteases of seeds from one species can act upon another when homogenized and co-incubated after germination. Among four different sprouts of chickpea, broccoli, lentil and mung we coincubated six pairwise combinations for the same time period as the homogenates alone were incubated (four hours). We identified the peptides from each source species separately, yielding 12 datasets; each of these had N-terminal and C-terminal datasets, whose proteolysis profiles can be compared to the four species homogenised but unmixed datasets.

Figures 4 and 5 present the proteolytic motif distributions for co-incubated samples, with rows representing the analysis of peptides matched to the row species, and columns representing the species with which that species was co-incubated. The diagonals represent the right hand column shown in figures 2 and 3, showing the incubated homogenate profiles of the four species on their own, repeated again to facilitate easier visual comparison.

**Figure 4.**
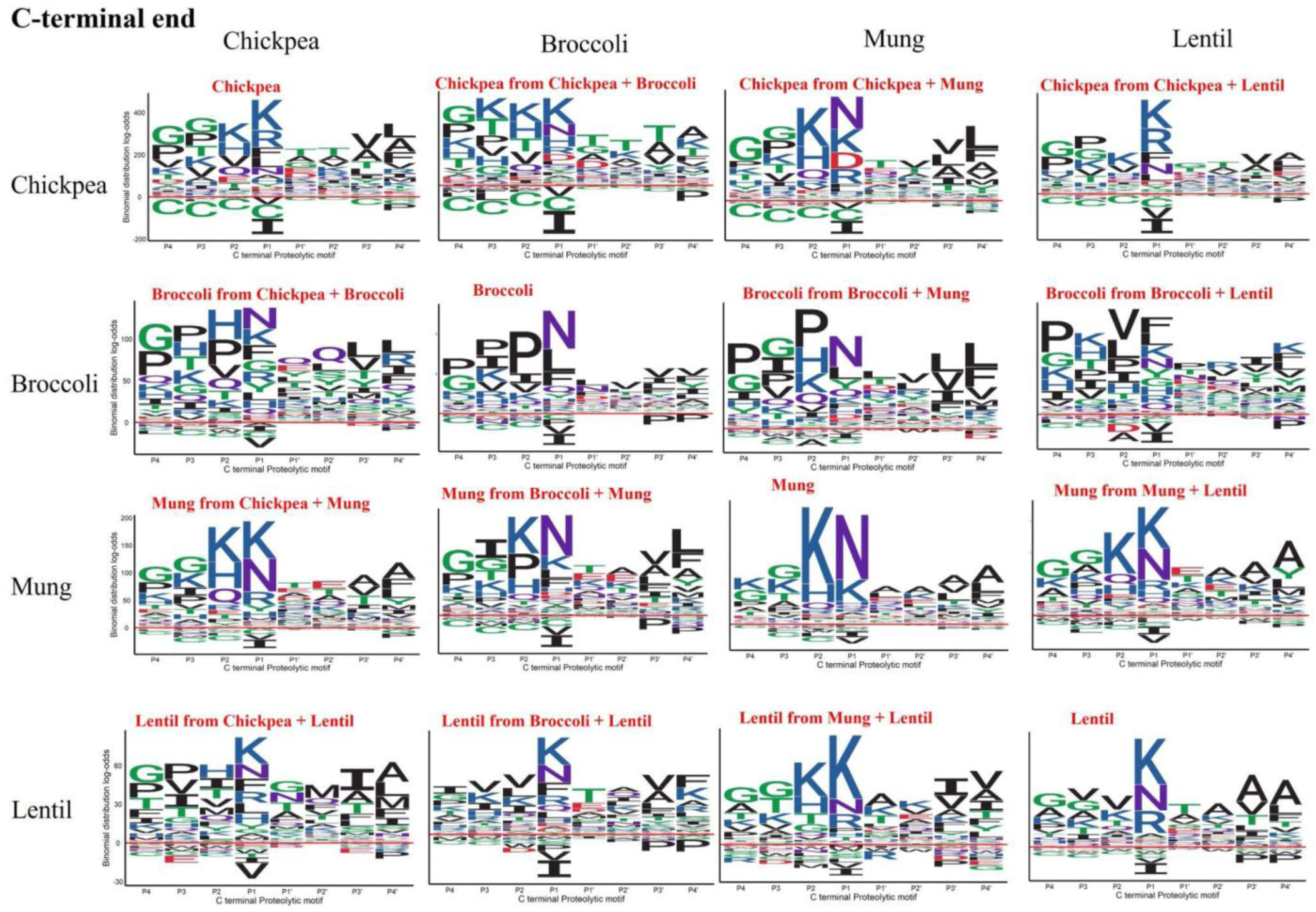
Proteolytic pattern of co-incubated sprouts at peptide C-termini. Rows: Profiles from the peptides assigned to that species. Columns: Species whose homogenate was coincubated with the species in the row.

**Figure 5.**
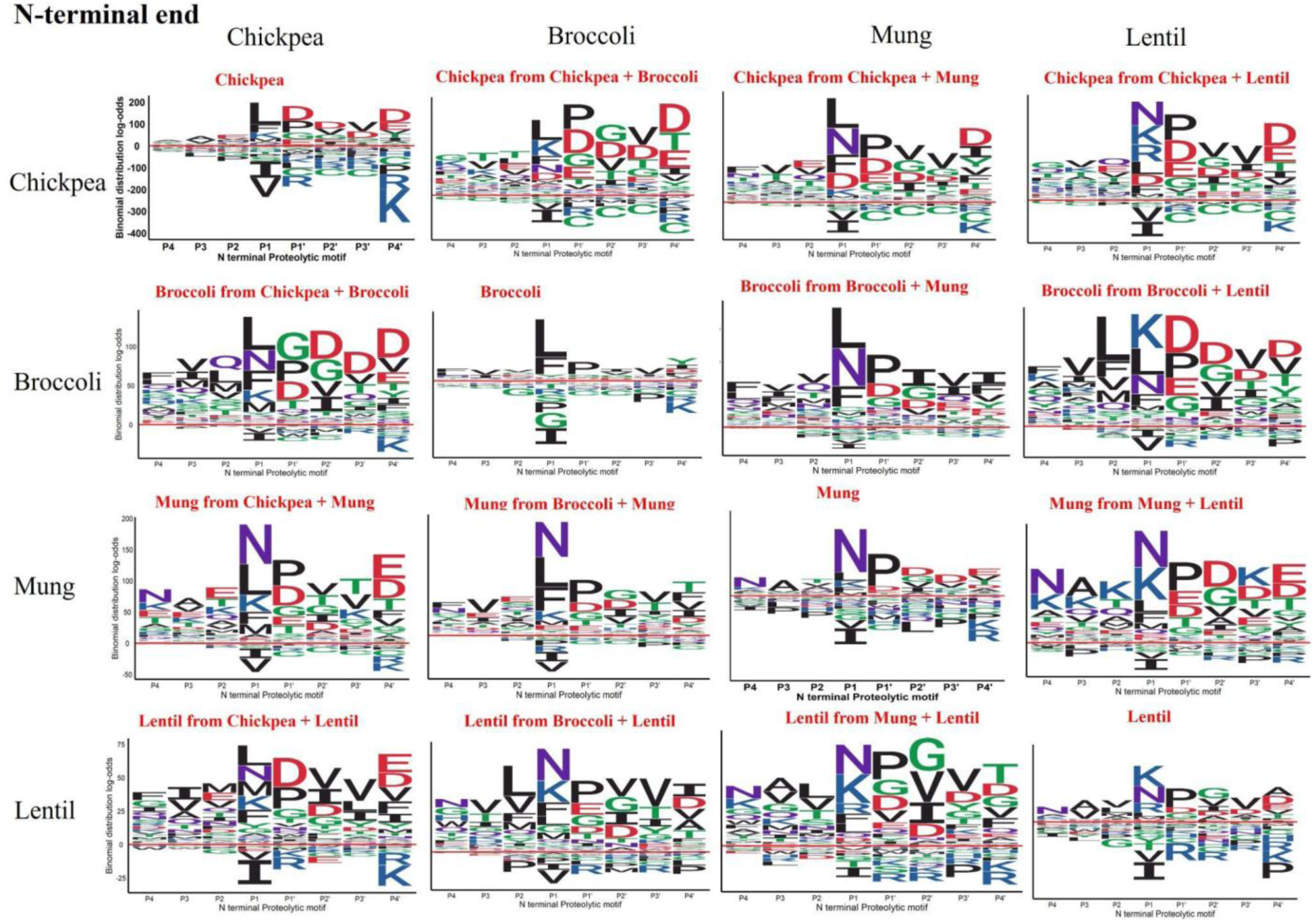
Proteolytic pattern of co-incubated sprouts at peptide N-termini. Rows: Profiles of peptides assigned to a species. Columns: Species whose homogenate was coincubated with the species in the row.

Since these data are complex to absorb visually, we have tabulated a summary of the most important differences, taking P4 to P1 preferences from the N-terminal dataset (Figure 5), and P1’ to P4’ preferences from the C-terminal dataset (Figure 4).

From Table 2 and Figure 4 and 5, it can be seen that co-incubation clearly enables an activity enriched in one species to act on the proteins of another species. Firstly, as might be expected, motif preferences that are associated with one species are seen to introduce novel cleavages at the termini of peptides of the co-incubated species (underlined in Table 2). Thus, both mung and lentil contribute a legumain-like cleavage at P1-N to chickpea. Lentil additionally contributes a P1-R cleavage of chickpea peptides in the mixture, suggesting a vignain-like cleavage distinct from the P1-K cleavage already present in chickpea. In addition, lentils contribute a vignain-like P1-K activity to cleavage of broccoli peptides. Lentil shows an increase in P1-F in the presence of broccoli. There is only one novel P1 preference that emerges that cannot be mapped to an activity not seen in one of the sprouts in isolation (italics in Table 2): chickpea shows a preference for caspase-like P1-D in the presence of mung. Thus, it appears in this case that a proteolytic activity absent in both species samples is activated on mixture, possibly through proteolytic activation of the protease, or through degradation of protease inhibitors. However, the mung peptides from the same sample do not show this presence, so the activity may be restricted to a particular class of cleavage substrates that are more common in chickpea than in mung. Together, these data provide evidence that the mixture of sprouts of different species supports cooperative proteolysis.

**Table 2.**
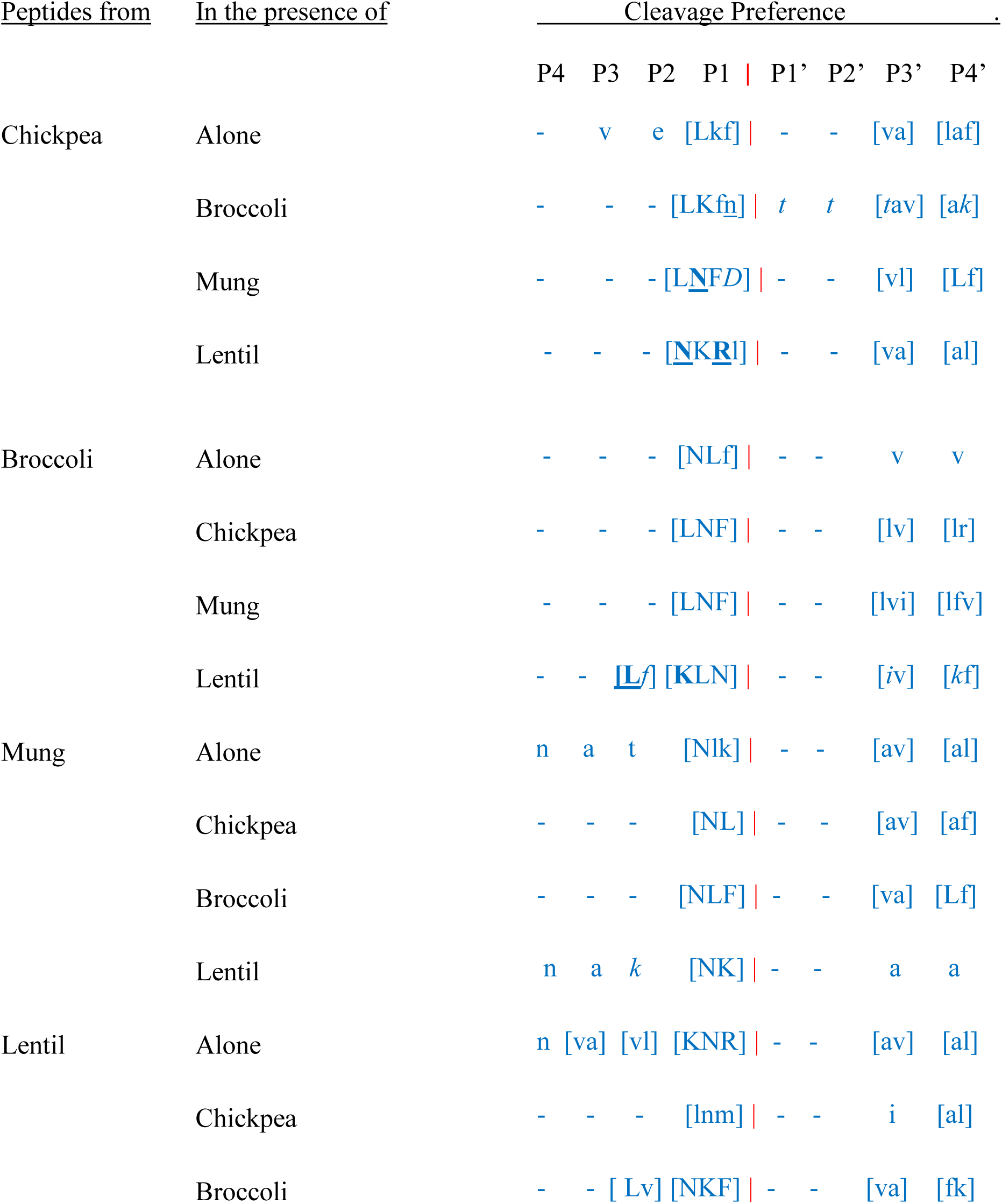
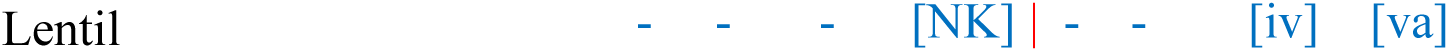
Summary of notable preferences. Amino acid preferences P4-P1 are from the N-termini, and P1’-P4’ from the C-termini. Red line: cleavage site. Square brackets: following standard regular expression [KH] means either K or H at that position; alternative amino acids are listed in order of how enriched they are, with less enriched amino acids in lower case. Underlined: amino acid preferences that appear to be more enriched from activity of co-incubated species (bold: more striking instances of this). Italic: amino acid preference not seen in either of the co-incubated species.

As discussed above, motifs accumulating at N or C termini of peptides during germination are likely to reflect a subset of persistent peptides that are difficult to break down further. It is of great interest whether the four apparently “persistent” peptide classes (negative charge near legume peptide N-termini, prolines near broccoli C-termini, G and K rich motifs at chickpea and mung C-termini, P1-P at N-termini of all four species) are reduced by mixture with any other species, which might be able to break them down. Proline remained consistently enriched at peptide N-termini (Figure 5) in all the species mixtures. There is some suggestion that lentils reduces the enrichment of G and K rich C-terminal motifs in chickpea peptides (figure 4).

To our surprise, the mixture with chickpea or lentil enriched the broccoli peptides for the acidic N-terminal motifs (rich in D and E) that are seen in legumes, but not in broccoli on its own. One possible explanation is that the broccoli aminopeptidase that effectively digests such peptides is down-regulated by chickpea and lentil inhibitors. Co-incubation with broccoli had little effect in eliminating these preferences from chickpea peptides (Fig. 5), and no marked effect in mung and lentils.

For “persistent” broccoli peptides with prolines at the carboxy terminal P4 to P2 positions (Fig. 2), addition of homogenates from the other three species somewhat altered this profile, it did not markedly reduce it (Fig. 4). This suggests that these proline rich peptides are not easily broken down by broccoli or legume peptides.

### 3.6 Quantification of species contributions to proteolysis

We used linear regression to quantify inter-species proteolytic effects. The goal was to capture not just the endoproteolytic preferences, but the sum of all effects, including endoproteolytic enrichment, rapid exoproteolytic removal of residues from termini and persistence of indigestible peptides. We analysed all eight positions of the proteolytic cleavage window. Given the diversity among N and C terminal patterns (partly reflecting likely strong differences in exoproteolytic patterns), we analysed them separately. Table 3 shows the results for C-terminal cleavage sites, with Table S1 showing those for N-termini. The regression analysis quantifies percentage contributions from each species in a mixture to the proteolytic profile of the peptides of each species, by making reference to their profiles seen in the single species homogenates alone. There was a tendency for peptides to be more likely to match the profile of their species of origin (Table 3), perhaps suggesting incomplete homogenisation of samples. There was statistical support for N-terminal contributions from both species (confidence intervals of the estimated contributions not overlapping 0 or 100%), for 9 of the 12 combinations, with mung peptides being apparently unaltered by mixture with lentil samples and vice versa, and broccoli not altering chickpea peptide profiles significantly (Table S1). For the C-terminal dataset, contribution of proteolytic activities from other species ranges from 5 to 58% (Table 3). Overall the estimated contributions from N and C-termini were similar, despite their markedly different profiles seen in Figure 4 compared to Figure 5; the 2 of the 12 differing findings with non-overlapping confidence intervals were for the impact of lentil mixture on mung peptides, and vice versa, where the N-terminal datasets assigned the proteolysis patterns to their own species (perhaps more dominated by endoproteolysis preferences), while the C-terminal datasets showed more contribution from the other species (likely to be more influenced by other effects). Given the broad similarity of findings from both termini, the overall conclusion that mixtures of species contribute significantly to the digestion of each other’s peptides appears to be well supported. Chickpea peptides may be inefficiently broken down by proteases from the other three species, since the estimated contribution ranged from 22% in mung to only 5% in broccoli. Chickpea has contributed most to the proteolysis of other species (Table 3).

**Table 3.** Contribution of proteases ( in % ) at C-termini on peptides of different species.

| SAMPLES | Contribution of proteolytic activities from |  |  |  |
| --- | --- | --- | --- | --- |
| Chickpea peptides | Chickpea | Broccoli | Mung | Lentil |
| Chickpea +Broccoli | 95±5 | 5±5 | — | — |
| Chickpea + Mung | 78±6* | — | 22±5* | — |
| Chickpea + Lentil | 84±11* | — | — | 16±10* |
| Broccoli peptides |  |  |  |  |
| Broccoli + Chickpea | 44±9* | 56±10* | — |  |
| Broccoli + Mung | — | 84±6* | 16±6* |  |
| Broccoli + Lentil | — | 78±10* | — | 22±10* |
| Lentil peptides |  |  |  |  |
| Lentil +Chickpea | 38±20* | — | — | 62±20* |
| Lentil + Broccoli | — | 37±8* | — | 63±7* |
| Lentil + Mung | — | — | 26±12* | 74±12* |
| Mung peptides |  |  |  |  |
| Mung +Chickpea | 63±7* | — | 37±7* | — |
| Mung + Broccoli | — | 58±6* | 42±6* | — |
| Mung + Lentil | — | — | 63±7* | 37±7* |

\* Significantly different from 0% and 100% (confidence intervals at 95%)

### 3.7 Changes in precursor proteins categories

High-confidence protein identifications (Protein Probability > 0.99) were sorted on peptide abundance and the top 100 proteins were selected for further analysis for each of the samples. Heatmaps of peptide counts per protein show that different timepoints and conditions show clear changes in the proteins contributing to released peptides in chickpea and broccoli (Figures 6 and 7). Proteins were manually assigned to categories, and those with sufficient numbers in a species were tabulated (Tables 4 to 7). For each time point, the relative abundance of each protein group was calculated as a percentage of the total protein intensity. Rows were scaled using z-score standardization. We tested whether the distribution of protein functional categories in peptides of a species changed significantly in germinated (e.g. 4 days) versus germinated+homogenised (e.g. 4.2 days), and between homogenised single species versus mixed species samples.

**Fig. 6.**
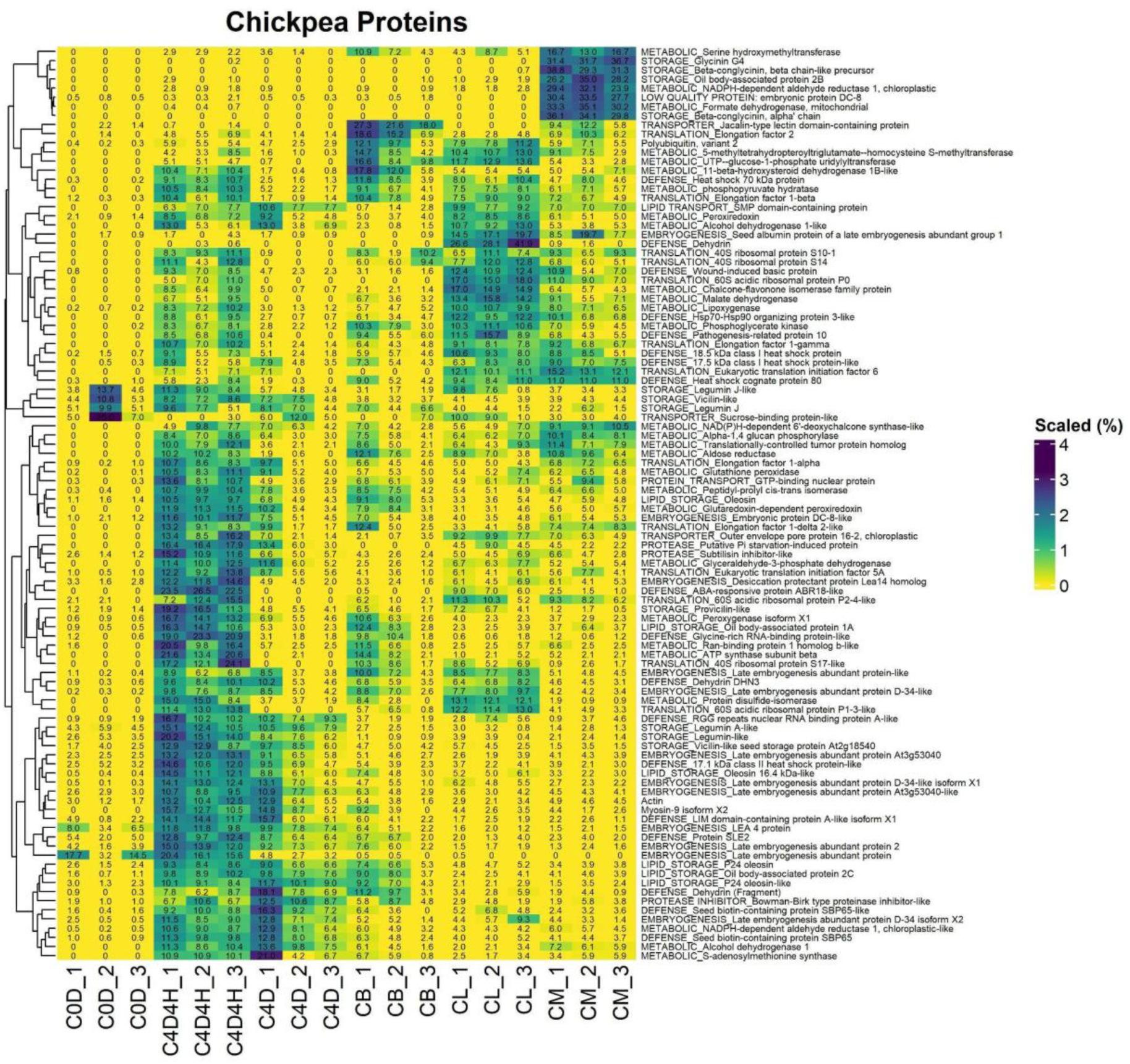
Heatmap chickpea precursor proteins calculated based on the number of peptides. Columns: C0D - soaked, C4D - germinated, C4D4H - incubated homogenate, CB - chickpea + broccoli, CL - chickpea + lentil, CM - chickpea + mung. The nos. in each cell represents the percentage of that particular protein (in rows) for that particular condition (in column).

**Fig. 7.**
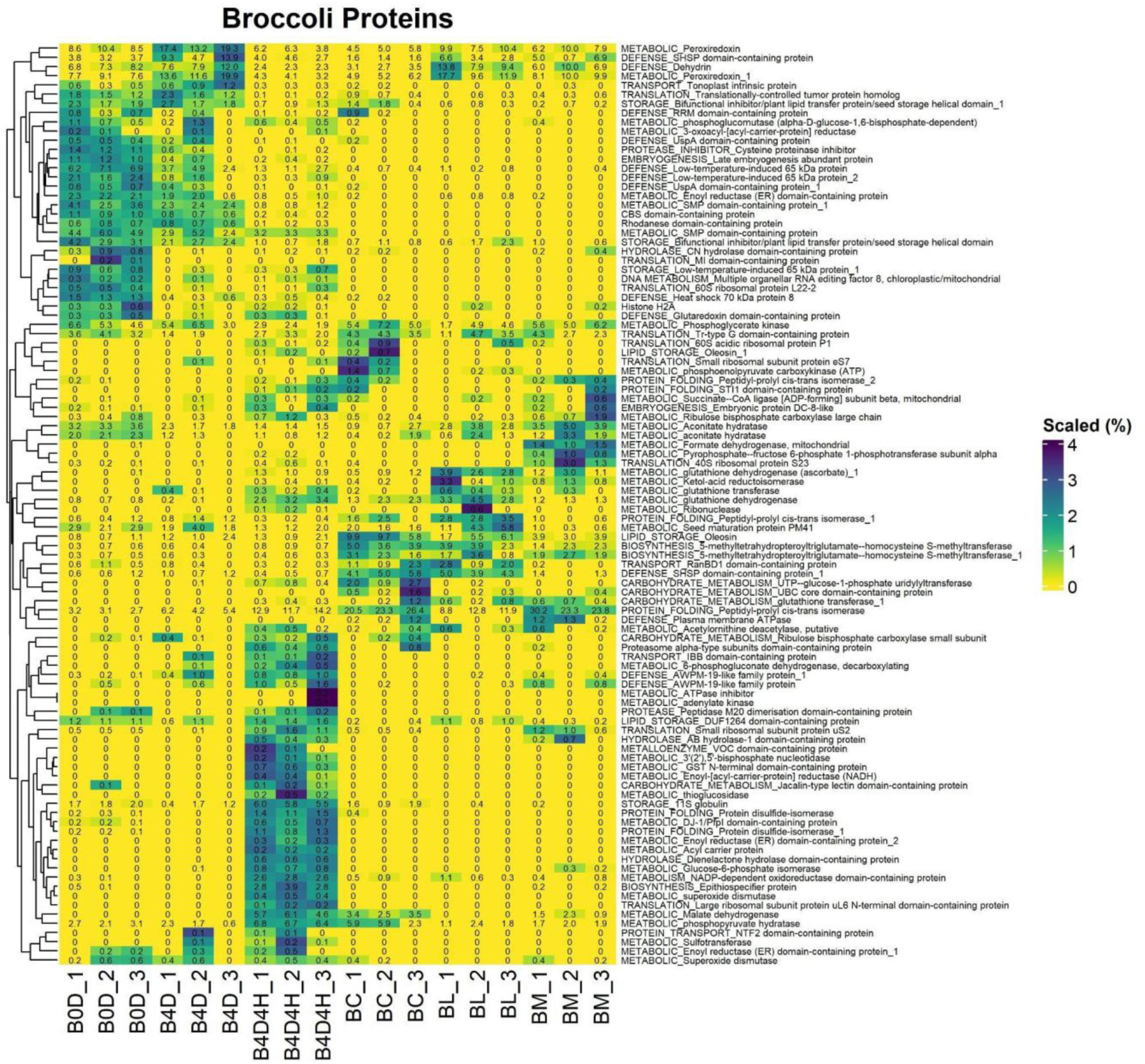
Heatmap broccoli precursor proteins calculated based on the number of peptides. Columns: B0D - soaked, B4D - germinated, B4D4H - incubated homogenate, BC - chickpea + broccoli, BL - broccoli + lentil, BM - broccoli + mung. The nos. in each cell represents the percentage of that particular protein (in rows) for that particular condition (in column).

**Table 4:** Average of % peptides counts in chickpea for different classes of proteins at different germination time points and statistically significant in incubated homogenate as compared to germinated sample (*) and in mixture of chickpea as compared to incubated homogenate (#)

| Family | Chickpea soaked | Chickpea germinated | Chickpea incubated homogenate | Chickpea +Broccoli | Chickpea + Lentil | Chickpea + Mung |
| --- | --- | --- | --- | --- | --- | --- |
| EMBRYOGENESIS | 28.38 | 16.42 | 13.70 | 9.911 | 10.41<sup>#</sup> | 6.90<sup>#</sup> |
| LIPID_STORAGE | 6.67 | 9.53 | 6.7943 | 8.39 | 3.60<sup>#</sup> | 3.26<sup>#</sup> |
| TRANSPORTER | 10.96 | 2.51 | 1.50 | 6.170 | 2.50 | 2.62 |
| METABOLIC | 3.19 | 23.30 | 26.58<sup>*</sup> | 31.09 | 30.84<sup>#</sup> | 31.55<sup>#</sup> |
| STORAGE | 27.9 | 10.80 | 8.76 | 5.36 | 4.99 | 21.04<sup>#</sup> |
| DEFENSE | 14.51 | 19.08 | 19.67 | 17.13 | 22.99 | 11.53<sup>#</sup> |
| TRANSLATION | 3.24 | 5.59 | 13.77<sup>*</sup> | 14.64 | 17.50 | 13.14 |

Homogenisation altered the protein origins of peptides from all four species, but with distinct differences. In chickpea, homogenisation significantly doubled the proportion of peptides derived from translation (5.6% to 13.8%) proteins (Table 4). After homogenisation, broccoli (Table 5) showed a striking rise in peptides from protein-folding proteins (5.3% to 14.4%) and a sharp decline in those from defence proteins (23.2% to 9.6%). In lentils (Table 6), homogenisation was marked by a fourfold increase in metabolic peptides (4.0% to 22.8%) and a doubling in peptides from defence proteins (11.6% to 23.4%). By contrast, mung homogenates showed a marked increase in peptides from lipid storage proteins, and a very sharp reduction in those derived from metabolic proteins (Table 7). Thus, the breakdown of tissue and cellular barriers by homogenisation appears to increase the access of certain proteins to novel proteolysis, in all four sprouts.

**Table 5:** Average of % peptides counts in broccoli for different classes of proteins at different germination time points and statistically significant in incubated homogenate as compared to germinated sample (*) and in mixture of broccoli as compared to incubated homogenate (#)

| Family | Broccoli soaked | Broccoli germinated | Broccoli incubated homogenate | Chickpea +Broccoli | Broccoli + Lentil | Broccoli + Mung |
| --- | --- | --- | --- | --- | --- | --- |
| BIOSYNTHESIS | 0.70 | 0.35 | 3.99* | 4.17 | 3.36 | 2.10 |
| PROTEIN_FOLDING | 3.19 | 5.26 | 14.36* | 23.56 | 11.17 | 25.81 |
| CARBOHYDRATE_METABOLISM | 0.13 | 0.23 | 1.05* | 2.84 | 0.20<sup>#</sup> | 0.12<sup>#</sup> |
| METABOLIC | 32.86 | 33.34 | 30.66 | 21.95 | 28.25 | 28.51 |
| STORAGE | 7.21 | 5.65 | 9.79 | 11.02 | 6.98 | 4.47<sup>#</sup> |
| DEFENSE | 20.62 | 23.25 | 9.60* | 6.24<sup>#</sup> | 15.39 | 13.37 |
| TRANSLATION | 6.34 | 2.98 | 4.83 | 5.88 | 3.55 | 6.21 |

**Table 6:** Average of % peptides counts in lentil for different classes of proteins at different germination time points and statistically significant in incubated homogenate as compared to germinated sample (*) and in mixture of lentils as compared to incubated homogenate (#)

| Family | Lentil soaked | Lentil germinated | Lentil incubated homogenate | Lentil +Broccoli | Lentil + Chickpea | Lentil + Mung |
| --- | --- | --- | --- | --- | --- | --- |
| EMBRYOGENESIS | 29.69 | 22.36 | 9.86 | 8.53 | 14.25<sup>#</sup> | 16.32<sup>#</sup> |
| LIPID_STORAGE | 6.24 | 11.94 | 0.60 | 0.08 | 7.06<sup>#</sup> | 6.52<sup>#</sup> |
| TRANSPORTER | 4.82 | 3.66 | 4.92 | 6.78 | 0.37<sup>#</sup> | 3.66 |
| METABOLIC | 1.37 | 3.96 | 22.75* | 46.26<sup>#</sup> | 34.71<sup>#</sup> | 27.22 |
| STORAGE | 28.17 | 35.65 | 20.02 | 10.67<sup>#</sup> | 6.34<sup>#</sup> | 26.37 |
| DEFENSE | 14.92 | 11.62 | 23.38* | 11.11<sup>#</sup> | 16.04<sup>#</sup> | 6.63<sup>#</sup> |
| TRANSLATION | 3.77 | 1.63 | 4.967 | 6.62 | 8.16<sup>#</sup> | 2.07<sup>#</sup> |

**Table 7:**
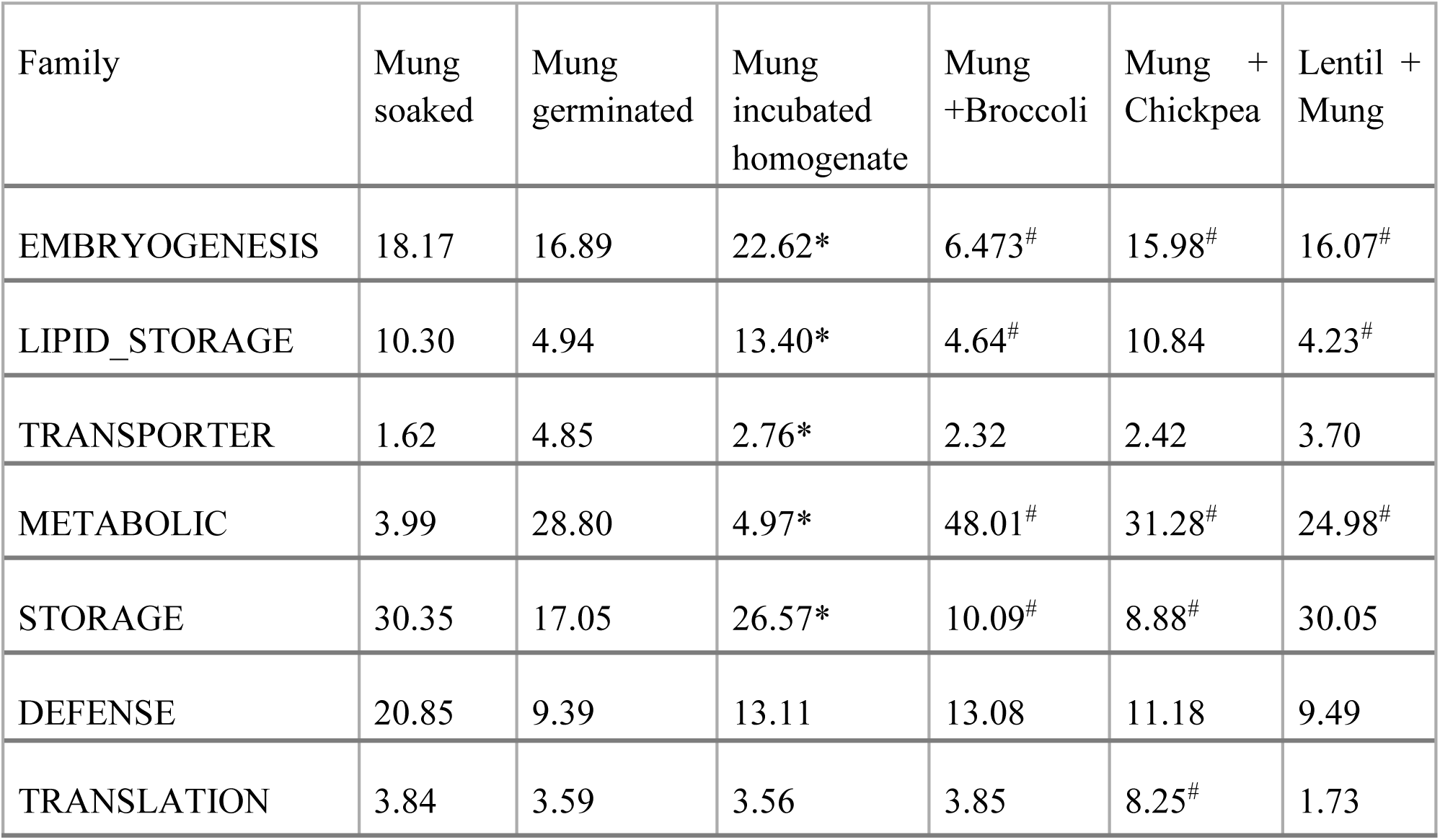
Average of % peptides counts in mung for different classes of proteins at different germination time points and statistically significant in incubated homogenate as compared to germinated sample (*) and in mixture of mung as compared to incubated homogenate (#)

When species were mixed, there were many further shifts in the categories of proteins releasing peptides (Tables 4-7). . While chickpea mixtures maintained high levels of chickpea peptides from metabolic proteins (31%), the most notable shift was a doubling of peptides from chickpea storage proteins by mixture with mung (Table 4). Lentil and mung both showed a very marked increase in metabolic protein-derived peptides, and a decrease of storage protein-derived peptides, on mixture with broccoli or chickpea homogenates (Tables 6 and 7). In the case of mung, this correlates with the dominating P1-L cleavage seen n chickpea and broccoli homogenates (Figure 3), reflected in an increase in P1-L cleavage in the mung peptides from mixtures with chickpea and broccoli (Figure 5). There was also an increased release of broccoli peptides from protein-folding proteins on mixture with chickpea and mung, which may relate to an increased legumain-like (P1-N) cleavage pattern increased among broccoli peptides from these mixtures (Figure 5). Curiously, this pattern is enriched in mung but not chickpea homogenates (Figure 5), suggesting some complex regulatory shift on mixture, such as for example a breakdown of legumain inhibitors.

Taken together, these results show that homogenisation accentuates species-specific proteolytic preferences, which are further modulated by interspecies mixing. Germination, homogenisation and mixing with other species each increased the release of peptides from chickpea and lentil metabolic proteins (Tables 4 and 6), and broccoli showed a similar trend towards protein-folding proteins (Table 5). While germination and species mixture increased metabolic protein peptide release from mung, homogenisation actually reduced it, with a matching increase in the proportion derived from storage proteins, and categories linked to proteins stored in the seed (lipid storage, embryogenesis and defence proteins; Table 7).

### 3.7 Conclusions

Proteolysis during plant germination is a highly regulated and dynamic process that responds to both environmental conditions and developmental cues. The impact of different stages and components of the proteolytic activation has a strong potential to impact on the health and pathogen resistance of the sprouting seed, as well as the nutritional and functional attributes of plant-derived foods. Understanding how these enzymes function, interact with substrates, and respond to biotic and abiotic stimuli is essential for leveraging their benefits in agriculture, food technology, and nutrition.

The four dicot species we investigated, the legumes mung, chickpea and lentil and the brassica broccoli, lay down their major seed storage proteins in the dicotyledon seed leaf structures, in contrast with monocot grains, where the main storage compartment is comprised of triploid endosperm. While the proteolytic preferences of all four species clearly differed from each-other in both soaked and germinated seeds, the major preferred activities in legumes were legumain-like P1-N, vignain-like P1-[RK], metacaspase-like P1-D, phytepsin-like P1-[LF], and an L-removing carboxypeptidase. Broccoli was more limited in its main cleavage preferences, showing no strong vignain-like or metacaspase-like activities; however, metacaspase activity in legumes appears to occur in a short time period, so it may well be that those activities will come to the fore at different timepoints outside of those observed here.

Homogenisation of sprouts appears to further activate proteolysis, detected here both as shifts in the proteolytic preferences at the termini of peptides, and as shifts in the functional categories of proteins that are being actively broken down. In summary, homogenisation appears to induce increased legumain activity in mung and lentils, to reduce metacaspase-like peptide release in lentils, and increase phytepsin-like activity in broccoli. Homogenisation does sometimes reduce classes of accumulating less digestible peptides, but may accelerate the accumulation of others, possibly by digesting background peptides faster such that the undigested peptides come to dominate the dataset more clearly. Thus, overall homogenisation does not necessarily create a universal “soup” of increased activity across all proteases, with some activities likely to go up, as substrate-enzyme interactions increase by bringing together new combinations of proteases and substrates from separate compartments, while others go down, as the optimal conditions for particular proteases are altered. Changes in activities thus reflect the complexity of the set of proteases, substrates and inhibitors present in compartments that come together during homogenisation. The magnitude of the differences seen (e.g. the marked replacement of legumain-like cleavage after asparagine in broccoli with cleavage after leucine or phenylalanine; the over two-fold increase in peptides digested from mung lipid storage proteins) suggest that homogenisation of sprouts is likely to markedly alter various functional properties of sprouted seeds.

Mixtures of sprouted seed homogenates showed clear evidence that the proteolytic systems of one species can have a marked influence on the digestion of peptides from another species. This is not surprising, given the clear differences in proteolytic profiles among the profiles of the different homogenates (chickpea and broccoli favouring cleavage after leucine; mung after asparagine; lentil after lysine).

Both homogenisation and species mixture of homogenates incubated at room temperature are, like seed germination, more “natural” than many food processing steps used to facilitate protein hydrolysis. As such, they are less likely to result in damage to the food material, and can be readily incorporated into food preparation steps, with a potential to facilitate improvements with respect to texture, taste, digestibility and allergenicity (Wilson and Wilson, 2015). Peptides ending in leucine tend to taste bitter (Ishibashi et 1987), so adjusting germinating sprout conditions to optimise leucyl carboxypeptidase activity may have big impacts on the taste of a preparation. In conclusion, the triple activation of seeds opens up new avenues of research into the functional and health properties of germinated seeds that may be acceptable to consumers.

## Acknowledgements

We acknowledge Darragh Flynn for helping us with initial discussion on industry scale sprout germination conditions. Darragh Flynn and Flynn & Flynn Global Trade Ltd., T/A The Happy Pear, A67 EC56 Wicklow, Ireland are acknowledged for providing us with the seeds for the experiments.

## CRediT authorship contribution statement

Conceptualization - D.C.S., M.O’S., I.B.; Data curation - I.B., C.S. ; Formal analysis - I.B., R.F.D., D.C.S.; Funding acquisition -D.C.S.; Methodology - I.B., M.O’S., D.C.S.; Project administration - D.C.S., M.O’S.; Supervision -M.O’S., D.C.S., J.C; Visualization - I.B., R.F.D., D.C.S.; Writing - original draft – I.B., R.F.D., D.C.S; Writing – review and editing - all authors.

