## Supplementary table 1 for "Peptidomics Mapping of Proteolysis Highlights Triple Activation of Sprouted Seeds by Germination, Homogenisation and Species Mixture"

**Table S1 Contribution of proteases ( in % ) at N-termini on peptides of different species**

| **Chickpea peptides** | | | | |
| --- | --- | --- | --- | --- |
| Samples | Chickpea proteases | Broccoli proteases | Mung proteases | Lentil proteases |
| Chickpea +Broccoli | 98±5 | 2±5 | — | — |
| Chickpea + Mung | 84±9* | — | 16±9* | — |
| Chickpea + Lentil | 81±9* | — | — | 19±8* |
| **Broccoli peptides** | | | | |
| Broccoli + Chickpea | 55±10* | 45±10* | — | |
| Broccoli + Mung | — | 95[80,96] | 5[-3,10] | |
| Broccoli + Lentil | — | 38±12* | — | 62±11* |
| **Lentil peptides** | | | | |
| Lentil +Chickpea | 36±16* | — | — | 64±16* |
| Lentil + Broccoli | — | 21±11* | — | 79±11* |
| Lentil + Mung | — | — | 5±16 | 95±16 |
| **Mung peptides** | | | | |
| Mung +Chickpea | 50±7* | — | 50±7* | — |
| Mung + Broccoli | — | 56±6* | 44±6* | — |
| Mung + Lentil | — | — | 95±11 | 5±11 |

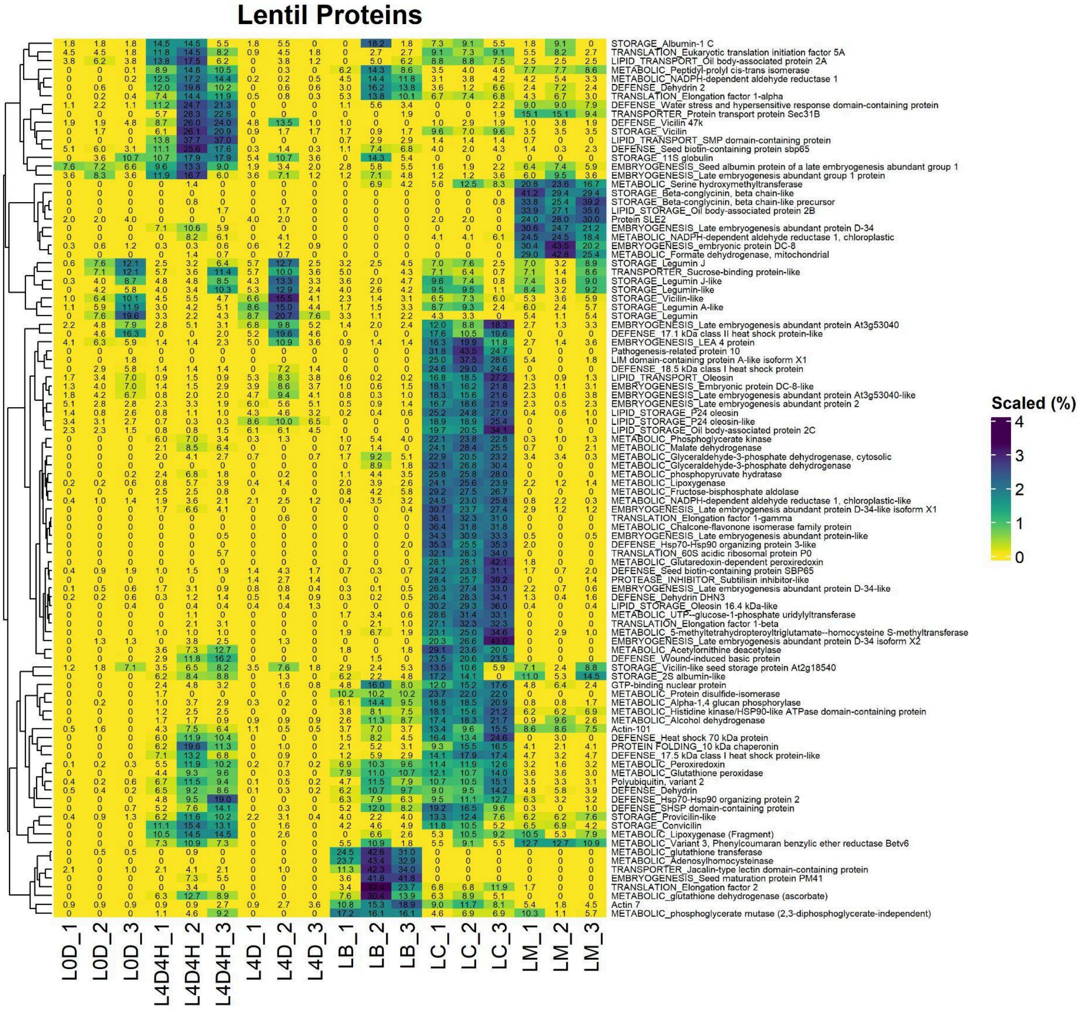


Fig. S1 Heatmap of lentil precursor proteins calculated based on the number of peptides. Columns: L0D - soaked, L4D - germinated, L4D4H - incubated homogenate, LC - chickpea + broccoli, LB - broccoli + lentil, LM - broccoli + mung. The nos. in each cell represents the percentage of that particular protein (in rows) for that particular condition (in column).

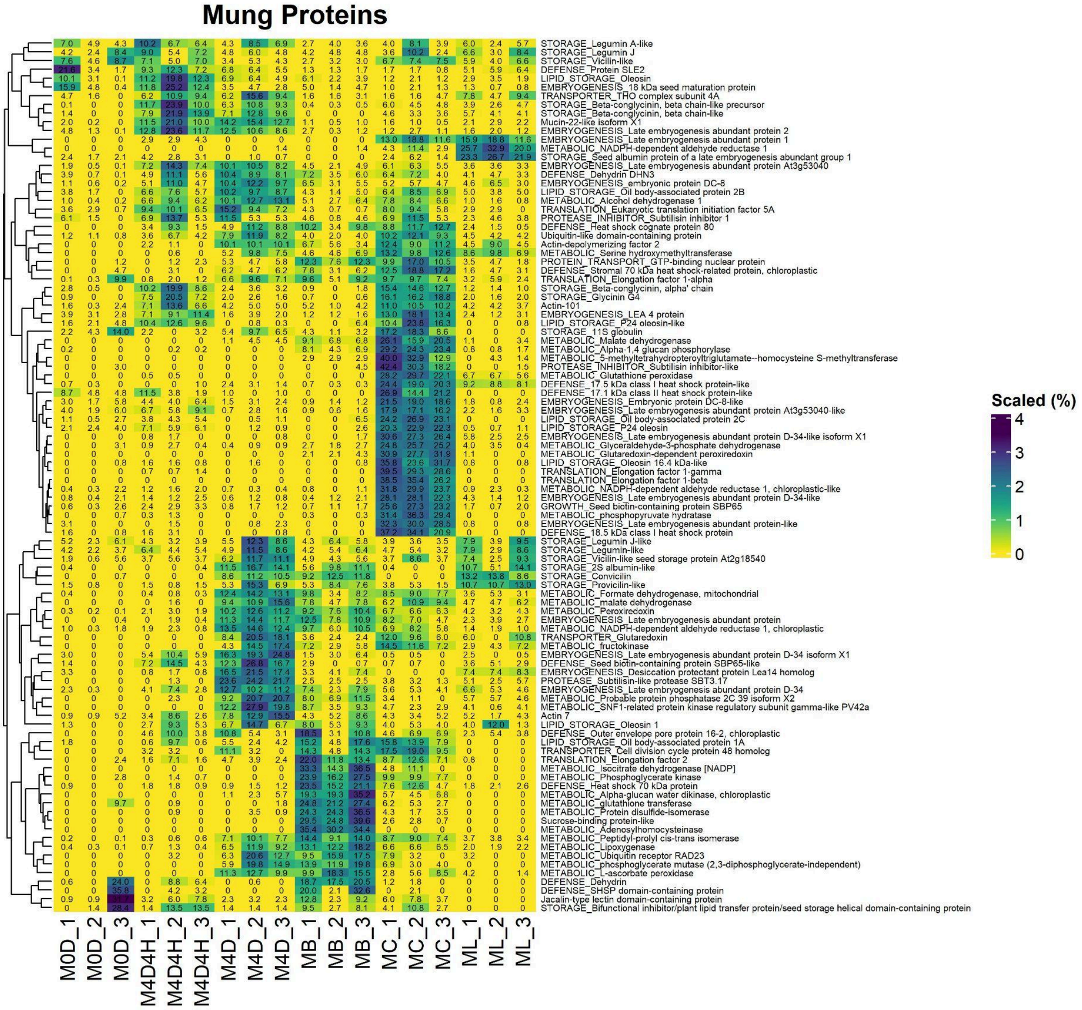


Fig. S2 Heatmap of mung precursor proteins calculated based on the number of peptides. Columns: M0D - soaked, M4D - germinated, M4D4H - incubated homogenate, MC - chickpea + broccoli, MB - broccoli + lentil, ML - broccoli + mung. The nos. in each cell represents the percentage of that particular protein (in rows) for that particular condition (in column).
